# Flow cytometry with cell sorting and sequencing as a tool for the study of the Humboldt Current krill stomach microbiota

**DOI:** 10.1101/2020.08.31.275677

**Authors:** Carlos Henriquez-Castillo, Belén Franco-Cisterna, Alejandro A. Murillo, Osvaldo Ulloa, Ramiro Riquelme-Bugueño

**Affiliations:** Instituto Milenio de Oceanografía (IMO), Universidad de Concepción, PO Box 160 C, Concepción, Chile; Departamento de Oceanografía, Facultad de Ciencias Naturales y Oceanográficas, Universidad de Concepción, PO Box 160 C, Concepción, Chile; Programa de Magýster en Ciencias mención Oceanografía, Facultad de Ciencias Naturales y Oceanográficas, Universidad de Concepción, PO Box 160 C, Concepción, Chile; Structural and Computational Biology Unit, European Molecular Biology Laboratory (EMBL), Heidelberg, Germany; Nordcee, Department of Biology, University of Southern Denmark, Odense, Denmark; Departamento de Zoología, Facultad de Ciencias Naturales y Oceanográficas, Universidad de Concepción, PO Box 160 C, Concepción, Chile

**Author notes:** **Corresponding Author:** Ramiro Riquelme-Bugueño, Departamento de Zoología, Facultad de Ciencias Naturales y Oceanográficas, Universidad de Concepción, PO Box 160 C, Concepción, Chile.

## Abstract

Euphausiids (or krill) are important contributors to marine biomass and key players in marine pelagic trophic webs. Euphausiids stomachs represent a specific niche for microbes that participate in the digestion of the host dietary components. To date, methods for the study of the diversity and function of these microorganisms remain complex. Often, bacterial ribosomal sequences obtained from lysates of stomachs are overrepresented by organisms from the surrounding environment. Flow cytometry with cell sorting (FC-CS) have become a powerful technique to study microbial community structure but also for the study of population genomics of gut-associated bacteria, even at a single-cell level.

In this study, we used FC-CS and sequencing of the bacterial 16S rRNA gene to study the microorganisms inhabiting the stomach of the Humboldt Current krill, *Euphausia mucronata*. This approach was complemented with DNA extraction and sequencing from whole lysate stomachs as described for other crustacean species.

Non-specific amplification was not retrieved in the polymerase chain reaction (PCR) from cells sorted, opposite to the observed using the DNA from the whole lysate. Sequences obtained from the whole stomach DNA were enriched in picocyanobacteria, meanwhile, sequences retrieved from cells sorted belonged almost exclusively to *Balneola* sp. of the new phylum, Balneolaeota. This study represents, to our knowledge, the first report of *Balneola* sp. in the stomach for any organism inhabiting the Humboldt Current System (HCS).

Our results suggest that the stomach-associated microbiota can be characterized by FC-CS and sequencing by manual scraping of the stomach coupled with the DNA extraction and sequencing. This work represents a baseline for similar studies of other mesozooplankton groups. The implementation of this technique might complement future studies on host-microbes’ interaction and their implications on the marine pelagic food web.

## INTRODUCTION

Krill are small zooplankton crustaceans found across the world’s oceans. Like other zooplankton groups, their stomachs represent a specific niche for diverse microorganisms which normally differ from the microbial communities in the surrounding seawater (*Tang et al., 2010*). Zooplankton bodies offer protection and an organic-rich micro-environment for the attached-bacteria (*Tang et al., 2010*), whereas bacteria can provide different metabolisms for maintaining the health of the host animals (*Shoemaker and Moisander, 2017*).

One of the most studied krill species is the Antarctic krill *Euphausia superba*, where bacterial growth occurs in the krill stomach’s, which is an important component of the entire digestive process of euphausiids. Bacterial growth in krill (*E. superba*) stomach has been suggested based on electron micrographs (*Rakusa-Suszczewski, 1988*). Also, diverse studies have been focused in the characterization of bacteria from krill stomach through different methodologies. These include: spread plate count method (*Kelly et al., 1978; Fevolden & Eidså, 1981; Donachie & Zdanowski, 1998*), acridine orange direct count under epifluorescence microscopy (*Fevolden & Eidså, 1981*), identification of cell sizes and morphology with optical and scanning electron microscopy (*Kawaguchi & Toda, 1997*), isolation and cultivation (*Donachie et al., 1995; Denner et al., 2001; Cui et al., 2016*), chromatographic analyses of bacterial proteins and enzymatic activity measurements with microassays (*Donachie et al., 1995*), and recently, the characterization of the bacterial diversity on the krill tissue has been done by using high-throughput sequencing (*Clarke et al., 2019*).

In the Humboldt Current System (HCS), one of the most productive marine systems in the world, the most abundant and endemic krill species is *Euphausia mucronata*. Its habitat is mainly restricted to the continental shelf in the coastal upwelling zones (*Riquelme-Bugueño et al., 2012*), where it has a high population density and biomass, contributing to the carbon cycling (*Gonzalez et al., 2009; Antezana, 2010; Riquelme-Bugueño et al., 2013*). These characteristics and their ecological role have recently prompted research in order to further our understanding about this species (*Gonzalez & Quiñones, 2002; Riquelme-Bugueño et al., 2015, 2016a,b*). Even though there has been quite a bit of progress in the study of krill ecology and physiology, little progress has been made in the last few years about the relationship between this krill species and their stomach-associated bacteria in the HCS, compared to the extensive analysis done for *E. superba* (Schmidt & Atkinson 2016).

To fill this gap and contribute to the study of *E. mucronata* stomach-microbiome, we used a complementary approach utilizing Flow cytometry and cell sorting (FC-CS) together with a conventional tissue DNA extraction, to assess the composition of the stomach microbial community. Flow cytometry is a high-precision technique that has been used intensively in microbiology since the early 1990s (*Amann et al., 1990*). It represents a specific approach for the counting of microbial cells. The cell-sorting capacity also enables further molecular analysis, allowing it to specifically characterize and quantify the microbiota component and predict its phylogenetic relationships. Flow cytometry and cell sorting (FC-CS) have become powerful techniques to study microbial community structure and population genomics of gut-associated bacteria at a single-cell level (*Koch et al., 2013; Engel et al., 2014*). In this work, we propose the use of Flow cytometry along with cell sorting to study the stomach-associated microorganisms of zooplankton species, in order to better understand the microbial composition of this largely unexplored ecological niche. We used the Humboldt Current Krill *E. mucronata*, as a study model in order to identify the microorganisms that are present in the krill stomach as well as their phylogeny. We also sought to explore new applications of flow cytometry in zooplankton ecology and the information that can be drawn from this technique.

## MATERIALS & METHODS

### Sampling

Sampling was carried out on March 2, 2016, at Station 18, located over the continental shelf off coast of central Chile (36.5°S, 73.1°W; seafloor 94 m depth) (Fig. S1). Physicochemical parameters were obtained using a conductivity-temperature-depth (CTD) SB911E profiler, equipped with an additional fluorescence sensor. Water samples were collected at night, using 10-L Niskin bottles on-board the R/V Kay Kay II (Department of Oceanography, University of Concepcion). The zooplankton was sampled from a depth of 50 m of the surface with a WP-2 standard plankton net (mesh size of 200 μm) and non-filtering cod ends. Once on board, live individuals were transported immediately to the laboratory for subsequent analyses. For flow cytometry analysis, of the planktonic free-living microbial community, 1.5 mL of seawater samples (in triplicates) were taken from 5, 10, 20, 50, 65, and 80 m depth, and were fixed on board with 10% dimethyl sulfoxide (DMSO) plus 0.05% of pluronic acid, maintained at room temperature for 20 min and quickly frozen in liquid nitrogen. The fixed samples were stored at −20° C until the analysis was performed.

### Krill stomach dissection and analyses

Specimens of *E. mucronata* were identified and separated (3 live individuals per analysis), washed with 0.2 μm filtered sterile seawater in sterile petri dishes under a laminar flow hood. Individuals were dissected, and their stomachs were extracted using sterile tweezers and scraped under a Stereo Discovery V8 zoom stereomicroscope (Zeiss). The stomachs were washed, dissected, and the content from three stomachs was pooled and re-suspended in 1 mL of sterile filtered seawater (Fig. 1) containing 10% of DMSO for flow cytometry analysis. Cell suspension was passed through a cell strainer (70-μm mesh) to remove particles that can clog the sample line of the flow cytometer and it was then stored in 1.5 mL sterile centrifuge tubes. Another set of three krill stomachs was dissected and stored in sterile 1.5 mL centrifuge tubes at −20° C for further DNA extraction. The genomic DNA was extracted from intact stomachs using the NucleoSpin Tissue XS kit (Macherey-Nagel®). The integrity of the DNA was checked in a 1% agarose gel, and the concentration was determined using a Qubit fluorometer V 1.27 (Invitrogen®). The DNA was then stored at −20° C until amplification.

**Figure 1:**
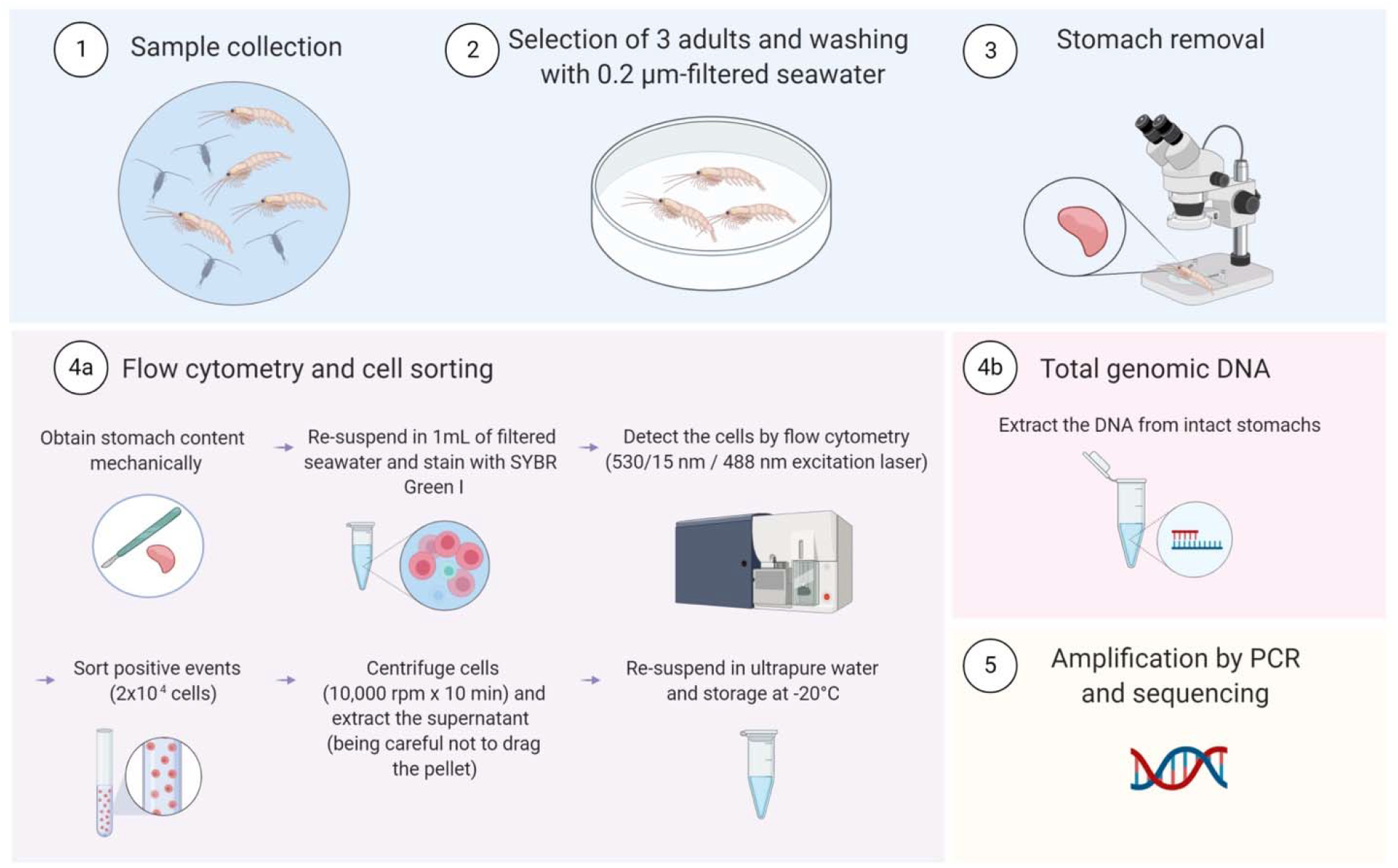
Methodology diagram. Schematic summary of the methodological procedures used in this study to identify microorganisms in the *Euphausia mucronata* stomach. The diagram was created using the BioRender (biorender.com) web platform.

### Flow cytometry and cell sorting analysis

Picoplankton in the water column was enumerated using a high-performance InFlux® flow cytometer (Becton Dickinson, formerly Cytopeia). Autofluorescent particles were identified by their red fluorescence, detected at 692/40 nm using three excitation lasers (457 nm, 488 nm, and 532 nm). For heterotrophic cells counts, samples were fixed with DMSO (10% final concentration), stained with the DNA dye SYBR Green I, as described in Marie et al. (*Marie et al., 1997*), and they were differentiated by light scatter (forward angle light scatter, FALS). SYBR Green I fluorescence was detected at 530/15 nm using a 488-nm excitation laser (Fig. 3). Each cytometer run was calibrated with 1 mm diameter fluorescent Ultra Rainbow beads (Spherotech Inc.). 100 μL for autofluorescent cells and 75 μL for heterotrophic cells were run at an average flow rate of 20 μL min^−1^ and monitored with a liquid flowmeter (Sensirion US). The events were recorded with Spigot software (Cytopeia), and FlowJo software v7.6.1 (Tree Star Inc.) was used for data analysis. Positive events for SYBR Green I fluorescence (530/20 nm) in stomach samples were sorted in the purity mode. The cytometer was configured in a two-tube mode, the sort chamber was UV sterilized, and 2 × 10^4^ cells from stomach samples were sorted into 1.5 mL sterile centrifuge tubes. Cells were centrifuged at 10,000 rpm for 10 min. The supernatant was removed, and the cells were re-suspended in 16.85 mL of ultrapure nuclease-free water (IDT technologies) and stored at −20°C until analysis (Fig. 3).

### Polymerase chain reaction (PCR) conditions

The 16S rRNA gene was amplified using the eubacterial 358F 5’-CCTACGGGAGGCAGCAG-3’ (*Muyzer et al., 1993*) and 907RM 5’-CCGTCAATTCMTTTGAGTTT-3’ (*Muyzer & Smalla, 1998*) primer pairs. The PCR amplifications were carried out with a total reaction volume of 25 μL per sample. Each mix contained 0.5 mM dNTPs, 0.75 mM MgCl_2_, 0.2 μM of each primer, 1 U of Taq polymerase, and 1X Go taq buffer (Kappa Biosystems, Wilmington, MA, USA). For total community PCR, 20 ng of stomach-extracted DNA was added. A mixture of the reagents was added directly into the tube containing the cells for the sorted samples. The amplification conditions consisted of initial denaturation at 95° C for 5 min, followed by 30 cycles at 94° C for 30 s, at 52° C for 30 s, and at 72° C for 1 min, with a final extension at 72° C for 10 min.

### Cloning, sequencing and phylogenetic analyses

The clone libraries were constructed from the PCR products, obtained from the DNA of both the whole-stomach microbial communities and the sorted populations, using the pGEM®-T easy Vector System (Promega). Duplicate PCR products from the samples were pooled and purified with the Wizard® Gel and PCR Clean-Up System (Promega), ligated and subsequently cloned into *E. coli* JM109 competent cells, following the manufacturer’s specifications (Promega). After a PCR screening was conducted, selected clones with the correct size were sequenced at Macrogen Inc. (Korea), and the sequences were deposited in GenBank under the accession numbers MG010923-MG011103.

The sequences retrieved in this work were quality filtered and then locally aligned against the latest Bacterial and Archaeal 16S rRNA database from the National Center for Biotechnology Information (NCBI), in addition to the IMG-ER annotated genomes using MEGABLAST. For Balneolales order phylogeny, operational taxonomic units (OTUs) having a sequence similarity of 97% and ones that matched Balneolales were aligned against reference sequences using SSU-Align. Phylogenetic reconstructions were performed on 560 aligned nucleotides. The phylogeny was inferred by the maximum likelihood (ML) method, using the generally time reversible parameter and assuming a discrete gamma distribution (GTR+G). The model was selected according to the Bayesian and Akaike information criterion, using the JModel test 2.1.3 (*Darriba et al., 2012*). The phylogenetic inference by ML was done with BOSQUE Software (*Ramirez-Flandes & Ulloa, 2008*). The topologies of the trees were obtained after ML analyses were done, and the robustness of inferred topologies were supported from 100 nonparametric bootstrap samplings for ML. The tree was drawn with iTOL (*Letunic & Bork, 2006*). To detect *Balneola*-related sequences from the study area, 16S rRNA bacterial sequences from the HCS were retrieved from NCBI (Accession numbers: KM461719, KM461941, DQ810296 and DQ810787.1; respectively n = 700). The sequences were locally aligned against our dataset by using MEGABLAST.

### Genomic potential of *Balneola* DSM 17893 and *Balneola* sp. EhC07

Cell motility, adhesion, and polymer degradation genes were searched in the available genome for *Balneola* DSM 17893 through the IMG-ER platform (https://img.jgi.doe.gov). Genes for *Balneola* sp. EhC07 were searched using the draft genome sequence, available under the GenBank accession number LXYG00000000.1.

## RESULTS

### Oceanographic setting

Photosynthetically active radiation (PAR) was 1.7 μE m^−2^ s^−1^ at surface strongly diminishing to 1 μE m^−2^ s^−1^ within the first 5 m of the water column. From this depth, PAR subtly decreased at a rate of 0.002 μE m^−2^ s^−1^ until 80 m. The above coincides with the peak of fluorescence, around 5 m depth, declining to cero at 20 m. Also, the oxygen concentration declined rapidly in the first 20 m of the water column, stablishing an oxygen minimum zone (OMZ) from 20 m until the sea bed (< 1 mL/L or ~20 μM of dissolved oxygen). The bacterial abundance presented two peaks, the first at 5 m depth, with a decline at the oxycline and a second peak around 65 m depth, in the core of the OMZ. The temperature varied from 13.5° C at the surface to 11.5° C at 80 m depth, with the thermocline (observed from 5 to 20 m) coinciding with the oxycline and showing a stratification of the water column (Fig. S1). This profile match with the upwelling conditions observed at that station during that season over time (*Sobarzo et al., 2007*).

### Flow cytometry analyses of the water column and krill stomachs

Different groups of autofluorescent picoplankton were observed, mainly in surface waters of the study area, including small photosynthetic eukaryotes and picocyanobacteria that differs in their pigment properties (Fig. 2A). These groups of autofluorescent organisms were not observed in the stomach samples (Fig. 2C), suggesting a niche differentiation between the water column and the krill stomach. In contrast, a clear fluorescent signature was observed in the stomach samples when stained with the DNA dye Sybr Green I (Fig. 2D), and this pattern differed in their optical properties to the patterns observed through the water column (Fig. 2B; Fig. S2). A specific aggregation of cells of similar optic characteristics was selected for cell sorting, indicated by the dotted line circle in figure 2, cytogram in panel D.

**Figure 2:**
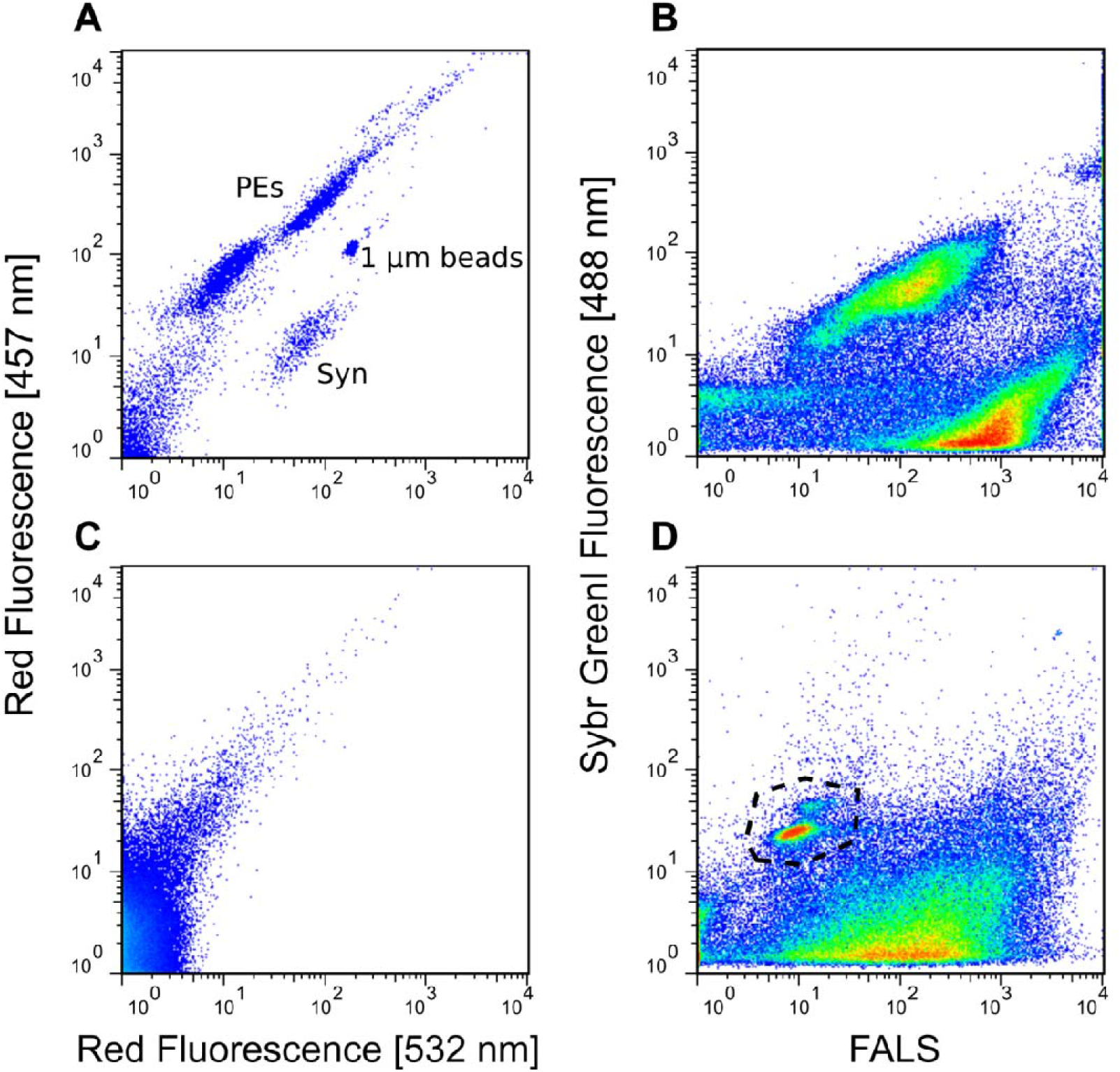
Flow cytometry plots from sorted seawater and *Euphausia mucronata* stomach samples collected from Station 18. Left panel: Detection of auto-fluorescent particles by using a two-laser approach. Red fluorescence was detected at 692 nm plus a 40 nm window, using a 532 nm green excitation laser (x axes) and a 457 nm blue laser (y axes). A: 5-meter depth seawater sample. C: *E. mucronata* stomach sample. Right panel: Detection of SYBR Green I fluorescence for picoplankton enumeration. Green fluorescence was detected at 530 nm plus a 15 nm window, using a 488-nm blue excitation laser (y axes). Forward angle light scatter (FALS) was used to identify particles. B: seawater sample at 5 meters. D: *E. mucronata* stomach sample. Syn: *Synechococcus*, PEs: Photosynthetic eukaryotes. The black dashed line in Fig. 1D indicates the sorted population for molecular analysis. The 1 μm beads are used as a size scale to identify the different populations of picoplankton.

### PCR amplification and sequencing of sorted cells and extracted DNA

Flow cytometry positive events for Sybr Green I fluorescence (dotted circle in Fig. 2D) were sorted and subjected to amplification of the bacterial 16S rRNA gene. The sorting procedure allowed for a specific amplification of the 16S rRNA gene fragment, contrary to the PCR products of multiple sizes obtained from the stomach-extracted DNA (Fig. S3). Most of the sequences retrieved from the amplification of the 16S rRNA gene, from the whole-stomach DNA extraction, were affiliated to picocyanobacteria (60.4%, n=52). Other genera found in this DNA sample, belonging to phylum Proteobacteria, where *Halioglobus*, *Tateyamaria*, *Sulfitobacter*, *Roseobacter*, and *Paracoccus* among others (Table 1). While sequences obtained from sorted stained cells from the pool of the three stomachs belonged almost exclusively to *Balneola* sp. (96.4%, n=80), a member of the recently defined phylum Balneolaeota. The closest cultured organisms corresponded to *Balneola* DG1502, a symbiont of the coccolithophore *Coccolithus pelagicus* f. *braarudii*, and *Balneola* sp. EhC07, a symbiont of the coccolithophore *Emiliania huxleyi* (Fig. 3), which were isolated from the South Pacific Ocean (*Green et al., 2015*).

**Figure 3:**
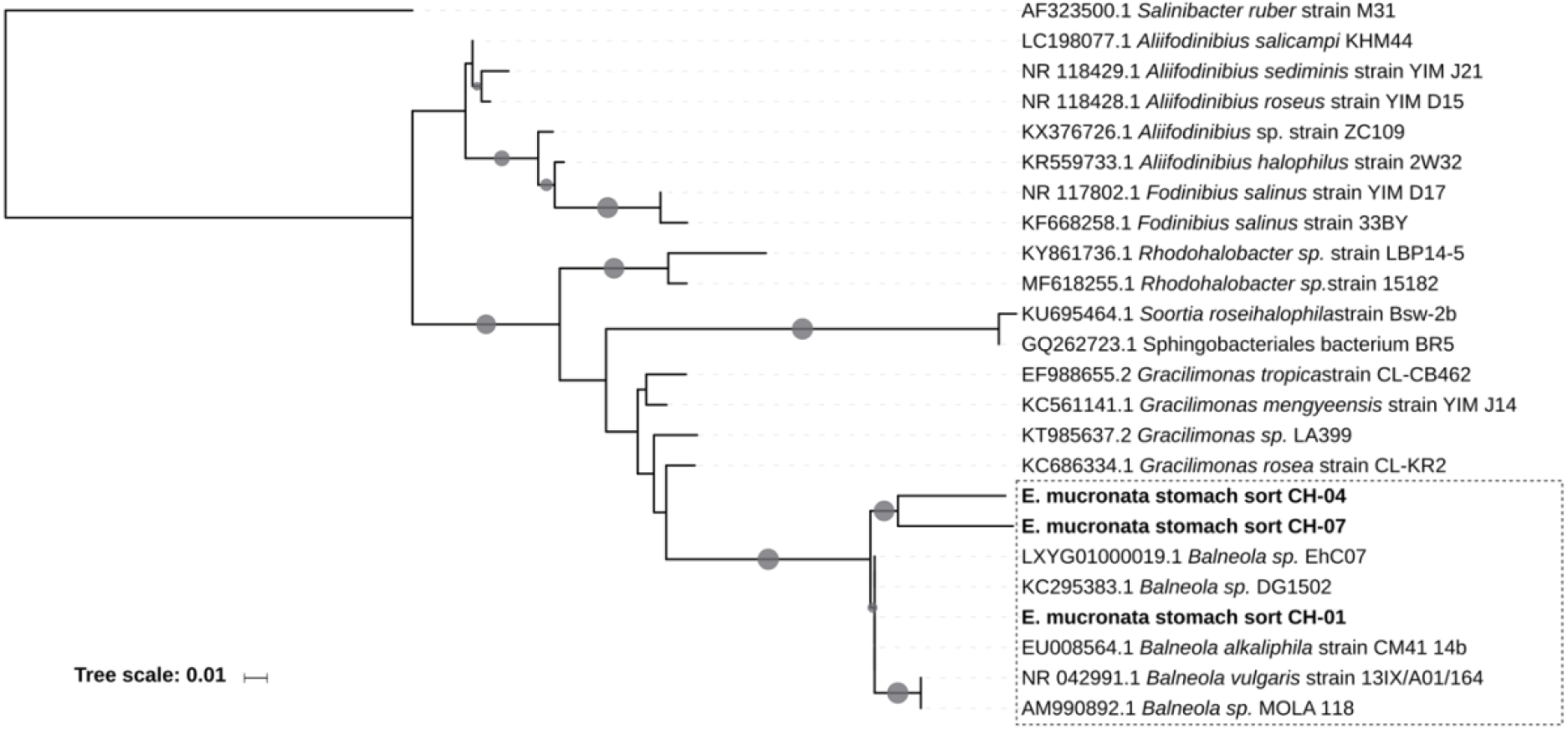
Phylogenetic tree (maximum likelihood) of 16S rRNA gene sequences obtained from sorted samples of *Euphausia mucronata* stomachs (dotted square), showing the affiliation within the family Balneolaceae. The tree includes the genus *Aliifodinibius*, *Balneola*, *Fodinibius*, *Gracilimonas*, and *Rhodohalobacter*. OTUs affiliated to *Balneola* were highlighted in bold. Bootstrap values of >50% are plotted at the nodes with grey circles. The size of the circles ranged between 50% and 100%. The tree-scale bar indicates the percentage of sequence divergence. *Salinibacter ruber* M31 was included as an outgroup.

**Table 1:** Phylogenetic affiliation, number, and percentage of sequences obtained from clones of sorted and DNA samples of the stomachs of *Euphausia mucronata* collected from Station 18.

| Affiliation<br>(**phylum**/genus) | N of sequences from |  | % of the total of sequences from |  |
| --- | --- | --- | --- | --- |
|  | Sorting | Total DNA from<br>stomach | Sorting | Total DNA from<br>stomach |
| **Proteobacteria** |  |  |  |  |
| *Pseudomonas* |  | 1 |  | 1.14 |
| *Vibrio* |  | 1 |  | 1.14 |
| *Lysobacter* |  | 1 |  | 1.14 |
| *Halioglobus* |  | 3 |  | 3.40 |
| *Luminiphilus* | 2 | 1 | 2.40 | 1.14 |
| *Haliaea* |  | 1 |  | 1.14 |
| *Lautropia* |  | 1 |  | 1.14 |
| *Tateyamaria* |  | 5 |  | 5.68 |
| *Sulfitobacter* |  | 3 |  | 3.40 |
| *Roseobacter* |  | 2 |  | 2.27 |
| *Paracoccus* |  | 2 |  | 2.27 |
| *Jannaschia* |  | 1 |  | 1.14 |
| *Citreimonas* |  | 1 |  | 1.14 |
| *Pseudoruegeria* |  | 1 |  | 1.14 |
| *Aliiroseovarius* |  | 1 |  | 1.14 |
| *Geoalkalibacter* |  | 2 |  | 2.27 |
| **Actinobacteria** |  |  |  |  |
| *Ilumatobacter* |  | 5 |  | 5.68 |
| *Cutibacterium* |  | 1 |  | 1.14 |
| **Firmicutes** |  |  |  |  |
| *Staphylococcus* |  | 1 |  | 1.14 |
| *Geobacillus* |  | 1 |  | 1.14 |
| **Balneolata** |  |  |  |  |
| *Balneola* | 80 |  | 96.38 |  |
| **Bacteroidetes** |  |  |  |  |
| *Polaribacter* | 1 |  | 1.20 |  |
| **Cyanobacteria** |  |  |  |  |
| *Synechococcus* |  | 18 |  | 20.45 |
| *Prochlorococcus* |  | 35 |  | 39.77 |

## DISCUSSION

The methodology used in this study, allowed us targeting specific bacterial populations from stomach samples based on their optical properties. Stomach-associated bacteria can be characterized by FC-CS and sequencing by manual scraping of the stomach, complemented with the DNA extraction and sequencing from the whole euphausiids’ stomachs. In this way, the optical properties of the stained cells indicated that sorted cells remained intact, inferred by the diameter measured with FALS parameter, and did not represent degraded DNA material, supporting the suitability of this methodology for an accurate analysis of the bacterial community from the stomach.

The intricate environment inside the krill stomach (Ullrich et al., 1991), probably fosters the development of unique niches for particular microorganisms. *Balneola* sp. EhC07 and *Balneola vulgaris* DSM 17893, whose genome information is available, contains a genetic repertoire for twitching (NCBI accession numbers: WP 018126390 and WP 066223579) and gliding motility (NCBI accession numbers: WP 018127305.1 and WP 066218663.1). These capacities favor physical contact of the bacteria with the host cells (*Tuson & Weibel, 2013*) and represent an advantage against the very intricate structure of euphausiids’ stomachs, as described for the Antarctic krill (*Ullrich et al., 1991*). Members of the order Balneolales have been found in different marine habitats. However, *Balneola* sp. have not been reported to be a free-living organism in the HCS (*Aldunate et al., 2018; Stevens & Ulloa, 2008*). Nevertheless, *Balneola alkaliphila* strain CM41_14b has been observed and isolated from surface waters, in the coastal north-western Mediterranean Sea (Urios *et al,* 2008). The functional capacities present in *Balneola* sp. for host colonization have also been reported in a wide range of plant and animal pathogens as well as in the formation of biofilms and fruiting bodies (*Green et al., 2015*; *Mattick, 2002*; *Rosana et al., 2016*). For example, *Balneola* sp. display the same capacities that are observed in *Polaribacter* spp., belonging to the phylum Bacteroidetes, which form an important part of the microbiota of marine organisms, especially in the gastrointestinal tract (*Moisander et al., 2015; Thomas et al., 2011*).

The lack of *Balneola*-related sequences in the whole stomach-extracted DNA may be related to the complexity of the euphausiids’ stomach (*Ullrich et al., 1991*) as well as methodological issues like the absence of a scraping of the stomachs prior the DNA extraction. It is possible that microorganisms were strongly attached to the stomach cells and cannot be disaggregated during extraction. Nevertheless, in the sorting approach, cell removal might be more efficient by scraping the stomach mucosa with a scalpel. This technique could facilitate cell recovery from the stomach tissue and, therefore, representing a technique improvement to study attached bacterial cells. The methodology used in this work allowed us to detect organisms that were not present in sequence libraries. Nevertheless, transcriptomic or stable isotopes analysis for the study of *in situ* activity are necessary to establish direct associations.

Microorganisms associated with the zooplankton digestive tract can be classified as resident, when bacteria are persistently present in the gut, and transient when bacteria do not form stable populations within the gut (*Tang et al., 2010*). In that sense, as cyanobacteria normally occur in the water column of the study area (*Iriarte et al., 2012*) and they are mainly grazed by nano and microplankton (*Böttjer & Morales, 2005*), its high abundance detected from the whole stomach-extracted DNA may represent transient microbiota passing through the digestive tract of krill during feeding. Whereas, *Balneola* sp. detected from sorted stained cells exhibit capacities favoring the physical contact with the host cells, therefore may be part of the resident bacteria associated with the Humboldt Current Krill. Nevertheless, experimental evidence is required to support this hypothesis.

The uses of different methodologies that can complement each other, such as flow cytometry with cell sorting coupled with whole stomach-extracted sequencing is proposed. The use of FC-CS can be also couple to high-throughput sequencing, to uncover the population diversity of groups of microorganisms with similar optic characteristics but with low abundance, and for single-cell genomics (SCG), isolating singular components of the community to explore its genetic repertoire. Both applications that will help to improve our understanding in future studies on host-microbes’ interactions and pelagic food webs.

## ACKNOWLEDGEMENTS

The authors would like to thank the captain and crew of Kay-Kay II research vessel (Department of Oceanography, University of Concepcion, Chile) for ongoing support at sea.

