## Supplementary material for "Flow cytometry with cell sorting and sequencing as a tool for the study of the Humboldt Current krill stomach microbiota"

Corresponding Author:

Ramiro Riquelme-Bugueño

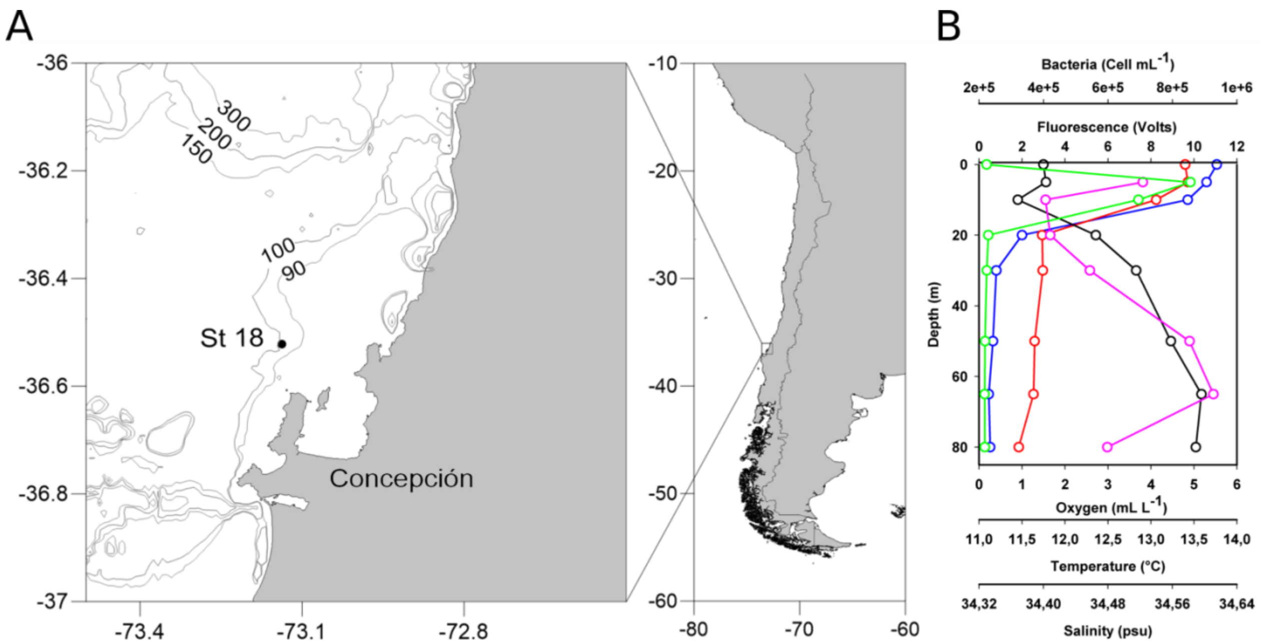


**Supplementary figure 1**: Area of study and samples collection. A shows the location of the oceanographic station 18 (St 18), off coast of Coliumo Bay, the seafloor at the station is at 90 m depth. B shows a vertical profile of physical-chemical parameters measured with a CTD, and the bacterial abundance from surface until 80 m depth, measured by flow cytometry. Color code: purple; bacterial abundance, red; temperature, blue; oxygen, green; fluorescence, black; salinity.

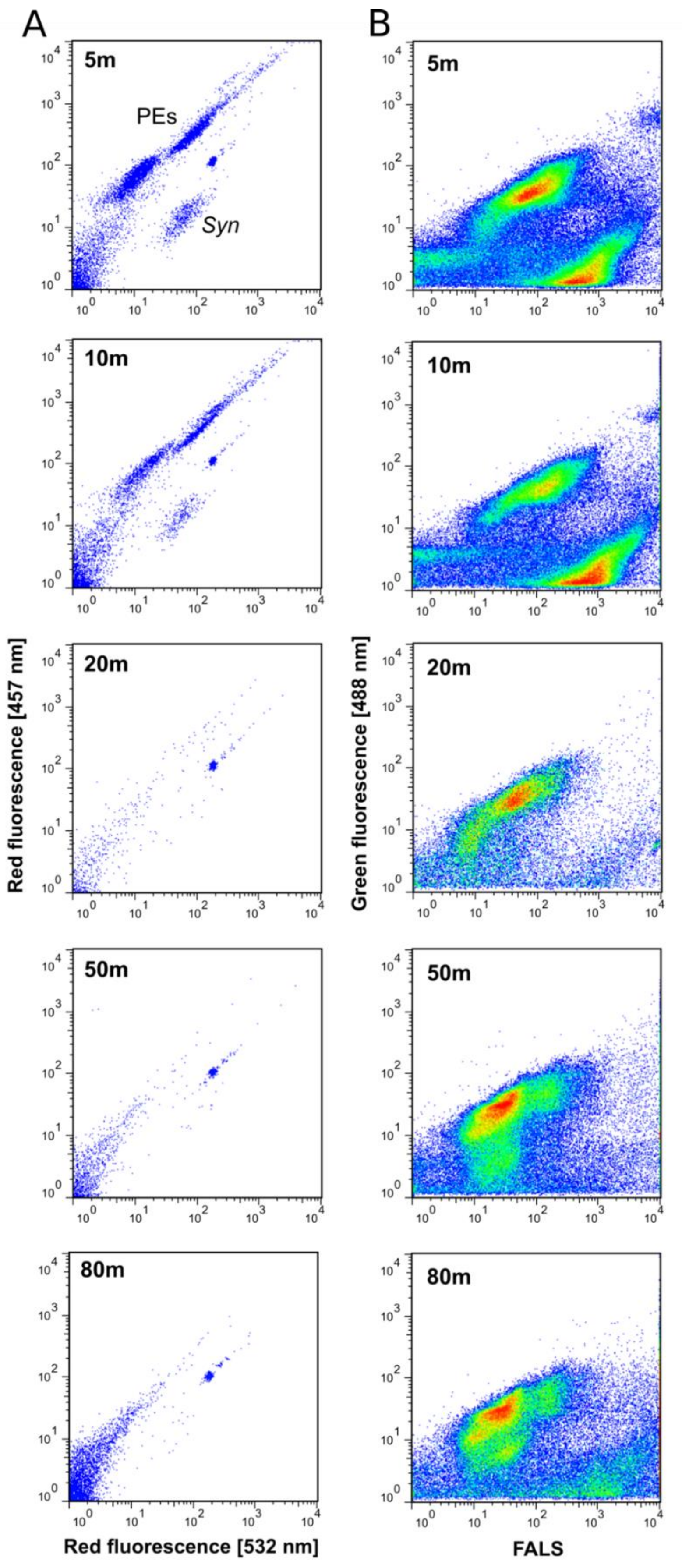


**Supplementary figure 2**: Vertical profile of flow cytometries for five depths in the water column of station 18. Cytograms in panel A show the auto-fluorescent organisms, through the detection of red fluorescence, where 2 populations are clearly distinguishable; Pico-eukaryotes (PEs) and the picocyanobacteria *Synechococcus* (*Syn*). The signal for these two populations practically disappeared at 20 m, confirming the results in Fig. S1B of fluorescence decay. Panel B shows the none auto-fluorescent organisms, stained with SYBR Green. The none-fluorescent bacterial community is distributed all along the water column, with a minimum at 20 m, coinciding with the oxycline.

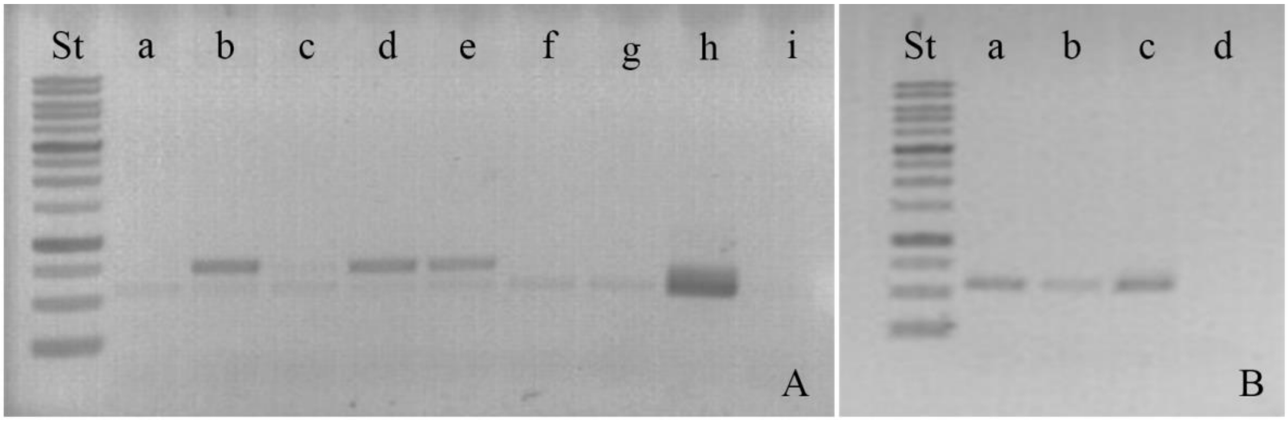


**Supplementary figure 3**: PCR amplification of the krill stomach content with primer pair 358F-907Rm. In A is presented the total community. Source of DNA for each lane: a) 4 stomachs, b) 7 guts, c) 2 stomachs, d) 5 guts, e) 10 guts, f) 1 stomach, g) 2 guts, h) positive, i) negative. In B are shown the sorted samples. Source of DNA for each lane: a) 2 x 104 sorted cells, b) 1.5 x 104 sorted cells, c) positive, d) negative.
